# Emerging genetic diversity of SARS-CoV-2 RNA dependent RNA polymerase (RdRp) alters its B-cell epitopes

**DOI:** 10.1101/2021.05.04.442686

**Authors:** Sushant Kumar, Gajendra Kumar Azad

**Author notes:** Corresponding Author: Gajendra Kumar Azad.

## Abstract

The RNA dependent RNA polymerase (RdRp) plays crucial role in virus life cycle by replicating the viral RNA genome. The SARS-CoV-2 is an RNA virus that rapidly spread worldwide and during this process acquired mutations. This study was carried out to identify mutations in RdRp as the SARS-CoV-2 spread in India. We compared the 668 RdRp sequences reported from India with the first reported RdRp sequence from Wuhan, China. Our data revealed that RdRp have acquired sixty mutations among Indian isolates. Our protein modelling study also revealed that several mutants including D833Y, A699S, Y149C and C464F can potentially alter stability and flexibility of RdRp. We also predicted the potential B cell epitopes contributed by RdRp and identified thirty-six linear continuous and twenty-five discontinuous epitopes. Among sixty RdRp mutants identified in this study, 40% of them localises in the B cell epitopes region. Altogether, this study highlights the need to identify and characterize the variations in RdRp to understand the impact of these mutations on SARS-CoV-2.

## INTRODUCTION

SARS-CoV-2 genome encodes 29 protein molecules which are categorised into three groups including structural, non-structural and accessory proteins (Gordon et al., 2020). SARS-CoV-2 has four structural proteins namely Spike glycoprotein, Membrane protein, Envelope protein and Nucleocapsid Phosphoprotein (Wu et al., 2020). It also encodes sixteen non-structural proteins (Nsp1-16) and nine accessory proteins (Chan et al., 2020). The non-structural proteins are involved in the maintenance of its genome, virulence and important steps of virus life cycle. The 16 non-structural proteins are synthesised as a single polypeptide molecule of 7096 amino acids known as Orf1ab that is subsequently cleaved into 16 separate proteins (Chan et al., 2020). The RNA dependent RNA polymerase (RdRp), also known as Nsp12, is a non-structural protein that replicates SARS-CoV-2 RNA genome (Wu et al., 2020). It associates with Nsp7 and Nsp8 and exist as a trimeric complex inside the viral envelope structure (Peng et al., 2020). By itself, RdRp has a very weak polymerase activity; however, the complex of RdRp with Nsp7 and Nsp8 significantly increases RdRp processivity and template affinity (Te Velthuis et al., 2012; Zhai et al., 2005).

RdRp of SARS-CoV-2 is 932 residues in length and contains distinct polymerase and nucleotide binding domains with a central connecting domain (Gao et al., 2020). Structurally, RdRp is comprised of an N-terminal β-hairpin (residues 31-50) followed by an extended nidovirus RdRp-associated nucleotidyl-transferase domain (NiRAN, residues 115-250) (Yin et al., 2020). Following the NiRAN domain is an interface domain (residues 251-365) connected to the RdRp domain (residues 366-920). Further, the domains of RdRp arranges in such a way that it forms a canonical right-handed cup configuration (Mcdonald, 2013), with the finger subdomain (resides 397-581 and residues 621-679) forming a closed circle with the thumb subdomain (residues 819-920)(Yin et al., 2020).

The SARS-CoV-2 was first reported from Wuhan province China in late 2019 (Wu et al., 2020). Wet sea food market area of Wuhan reported patients with pneumonia like symptoms, which was later identified as SARS-CoV-2 because of its similarity with SARS-CoV (Gorbalenya et al., 2020). The SARS-CoV-2 is highly contagious and causes mild to severe respiratory distress in infected individuals and the disease has been named as COVID-19 (World Health Organization, 2020). The SARS-CoV-2 spread very fast and within few months reached almost all countries around the globe (Worldometers, 2020). As of 02^nd^ May 2021, there are more than 150 million confirmed cases of COVID-19 and approximately 3 million deaths worldwide. This virus has caused global healthcare emergency because of its fast spread and sudden exponential rise in the COVID-19 patients and declared pandemic by world health organisation (WHO) (Cucinotta and Vanelli, 2020). This pandemic has caused enormous economic losses due to closure of most of the economic activities worldwide.

As the SARS-CoV-2 spread to other geographical areas with different climatic conditions compared to Wuhan, China, it started to mutate (Pachetti et al., 2020). The mutations acquired by the SARS-CoV-2 are retained as a consequence of natural selection, if the variants are more adaptable to the new conditions. In order to understand the variations occurring in RdRp among Indian geographical area, we analysed 668 RdRp sequences reported from India to identified sixty mutations. The B cell epitopes contributed by RdRp were predicted in silico, their functional consequences as well as the possible effect of mutations on them have been discussed.

## MATERIAL AND METHODS

### Protein Sequences retrieval from NCBI-virus-database

NCBI-virus-database is a repository for the nucleotide and protein sequences. This database also has protein sequences of SARS-CoV-2 that are available for users. We download the sequences of Orf1ab that contains all non-structural proteins including the RdRp sequence as described earlier (Azad, 2020). As of 10^th^ April 2021, NCBI-virus-database has 668 sequences deposited from the Indian COVID-19 patients. The supplementary table 1 shows the list of protein identifier accession number used in this analysis. The RdRp is located from 4393 to 5324 residues in Orf1ab which corresponds to 932 amino acids in length. The RdRp reference or wild type sequence was also downloaded from NCBI virus database. The accession number of reference sequence used in this study is YP_009724389 which is the first reported sequence of SARS-CoV-2 from Wuhan, China (Wu et al., 2020).

### Identification of RdRp mutants by multiple sequence alignments (MSAs)

In order to identify the variations present in the RdRp sequences among Indian isolates of SARS-CoV-2, the MSAs were conducted by Clustal omega programe (Madeira et al., 2019) as described earlier (Azad, 2021a). First, we uploaded the RdRp sequences in Clustal omega webserver and run the programe that uses HMM and pairwise alignment to generate the data. The variations from the reference sequence were properly marked and used for further analysis.

### B cell epitope prediction

B cell prediction methods to predict Linear B cell epitopes based on sequence characteristics of the antigen using amino acid scales and HMMs. The prediction of linear continuous B cell epitopes were conducted by IEDB (Immune Epitope Database and Analysis Resource) (Vita et al., 2019). The IEDB is an online server tool which predicts epitopes by a prediction method known as ‘Bepipred linear epitope prediction method 2.0’. For this prediction the threshold value of 0.500 was used during the evaluation.

The prediction of discontinuous B cell epitopes was performed by an online tool ‘DiscoTope 2.0’. For this prediction the threshold value was set at −3.7.

### Protein modelling studies

We performed protein modelling studies to understand the variation observed in the secondary structure might have any consequences on the three-dimensional structure of protein. This analysis was performed by DynaMut programe (Rodrigues et al., 2018) as described earlier (Azad, 2021b). This server uses the solved crystal structure of proteins and predicts the effect of mutation on the stability, intramolecular interaction and molecular fluctuations in the protein structure. For this study, we used recently reported structure of RdRp (PDB ID: 7BV1)(Yin et al., 2020). The effect of mutations on protein is shown in terms of difference in free energy (ΔΔG). The positive value indicates stabilizing mutation; however, negative value represents destabilizing mutation. DynaMut provides difference in vibrational entropy (ΔΔSvib ENCOM) between the wild type and mutant protein. The positive ΔΔSvib ENCOM indicates increase in protein flexibility and negative ΔΔSvib ENCOM represents increase in rigidity of protein structure due to mutations. We ran DynaMut webserver to calculate the ΔΔG and ΔΔSvib ENCOM that provides the impact of mutation on protein structure and stability. DynaMut server also provides the visual representation of the intramolecular interactions contributed by wild type and mutant residues with the neighbouring residues.

## RESULTS

### RdRp is frequently mutated among Indian isolates of SARS-CoV-2

In order to identify the mutations in RdRp, we compared the first reported sequence of RdRp from Wuhan, China with the sequences reported from India. In this analysis, we used 668 RdRp sequences reported from India and performed Clustal Omega mediated multiple sequence alignment with an aim to identify variations in amino acids between the sequences. The RdRp polypeptide sequence reported from Wuhan, China was used as wild type sequence. Our data revealed sixty mutations present among the Indian sequences of RdRp as shown in table 1. The table also shows the location of each mutation in the RdRp polypeptide sequences and the effect of mutation on amino acid charge and polarity. The mutations are also demonstrated on the schematic representation of RdRp as shown in figure 1A and B. Our result show that the mutations are spreading all over the RdRp polypeptide sequence. The distribution of mutations in different domains of RdRp has been highlighted in figure 1B. Our MSA data strongly indicates that RdRp is one of the most frequently mutated protein of SARS-CoV-2 because we observed sixty mutations in just 668 sequences analysed in this study.

**Table 1.** The list demonstrates the location and details of mutations of RdRp identified by Clustal Omega multiple sequence alignments. The RdRp sequence reported from Wuhan, China was used as wild type sequence for this analysis. The 668 sequences of RdRp reported from India were used for identifying mutations.

| S. No. | Wild-type | Charge Changes | Polarity Changes |
| --- | --- | --- | --- |
| 1 | K59N | Basic to Neutral | P to P |
| 2 | T85I | Neutral to Neutral | P to NP |
| 3 | K91N | Basic to Neutral | P to P |
| 4 | P94S | Neutral to Neutral | NP to P |
| 5 | A95V | Neutral to Neutral | NP to P |
| 6 | A97V | Neutral to Neutral | NP to P |
| 7 | I101L | Neutral to Neutral | NP to NP |
| 8 | M110K | Neutral to Basic | NP to P |
| 9 | R118C | Basic to Neutral | P to P |
| 10 | Y122F | Neutral to Neutral | P to NP |
| 11 | D140G | Acidic to Neutral | P to NP |
| 12 | T141I | Neutral to Neutral | P to NP |
| 13 | E144K | Acidic to Basic | P to P |
| 14 | T148I | Neutral to Neutral | P to NP |
| 15 | Y149C | Neutral to Neutral | P to P |
| 16 | R173H | Basic to Basic | P to P |
| 17 | A185V | Neutral to Neutral | NP to NP |
| 18 | M196I | Neutral to Neutral | NP to NP |
| 19 | I201L | Neutral to Neutral | NP to NP |
| 20 | T248I | Neutral to Neutral | P to NP |
| 21 | D269N | Acidic to Neutral | P to P |
| 22 | E278D | Acidic to Acidic | P to P |
| 23 | D284Y | Acidic to Neutral | P to P |
| 24 | T293I | Neutral to Neutral | P to NP |
| 25 | H295Y | Basic to Neutral | P to P |
| 26 | P323L | Neutral to Neutral | NP to NP |
| 27 | L329I | Neutral to Neutral | NP to NP |
| 28 | V342A | Neutral to Neutral | P to NP |
| 29 | V342L | Neutral to Neutral | NP to NP |
| 30 | V354A | Neutral to Neutral | P to NP |
| 31 | V354L | Neutral to Neutral | NP to NP |
| 32 | V398L | Neutral to Neutral | NP to NP |
| 33 | A406V | Neutral to Neutral | NP to NP |
| 34 | M463I | Neutral to Neutral | NP to NP |
| 35 | C464F | Acidic to Neutral | P to NP |
| 36 | I488S | Neutral to Neutral | NP to P |
| 37 | A526V | Neutral to Neutral | NP to NP |
| 38 | I536T | Neutral to Neutral | NP to P |
| 39 | V605A | Neutral to Neutral | NP to NP |
| 40 | M608I | Neutral to Neutral | NP to NP |
| 41 | L638F | Neutral to Neutral | NP to NP |
| 42 | T643I | Neutral to Neutral | P to NP |
| 43 | T644M | Neutral to Neutral | P to NP |
| 44 | S647I | Neutral to Neutral | P to NP |
| 45 | M668I | Neutral to Neutral | NP to NP |
| 46 | A699S | Neutral to Neutral | NP to P |
| 47 | L775P | Neutral to Neutral | NP to NP |
| 48 | E802A | Acidic to Neutral | P to NP |
| 49 | S814P | Neutral to Neutral | P to NP |
| 50 | Q822H | Neutral to Basic | P to P |
| 51 | D833Y | Acidic to Neutral | P to P |
| 52 | T848I | Neutral to Neutral | P to NP |
| 53 | S853L | Neutral to Neutral | P to NP |
| 54 | V880I | Neutral to Neutral | NP to NP |
| 55 | D893Y | Acidic to Neutral | P to P |
| 56 | Y908I | Neutral to Neutral | P to NP |
| 57 | T909I | Neutral to Neutral | P to NP |
| 58 | S913L | Neutral to Neutral | P to NP |
| 59 | S919L | Neutral to Neutral | P to NP |
| 60 | P925S | Neutral to Neutral | NP to P |

**Figure 1:**
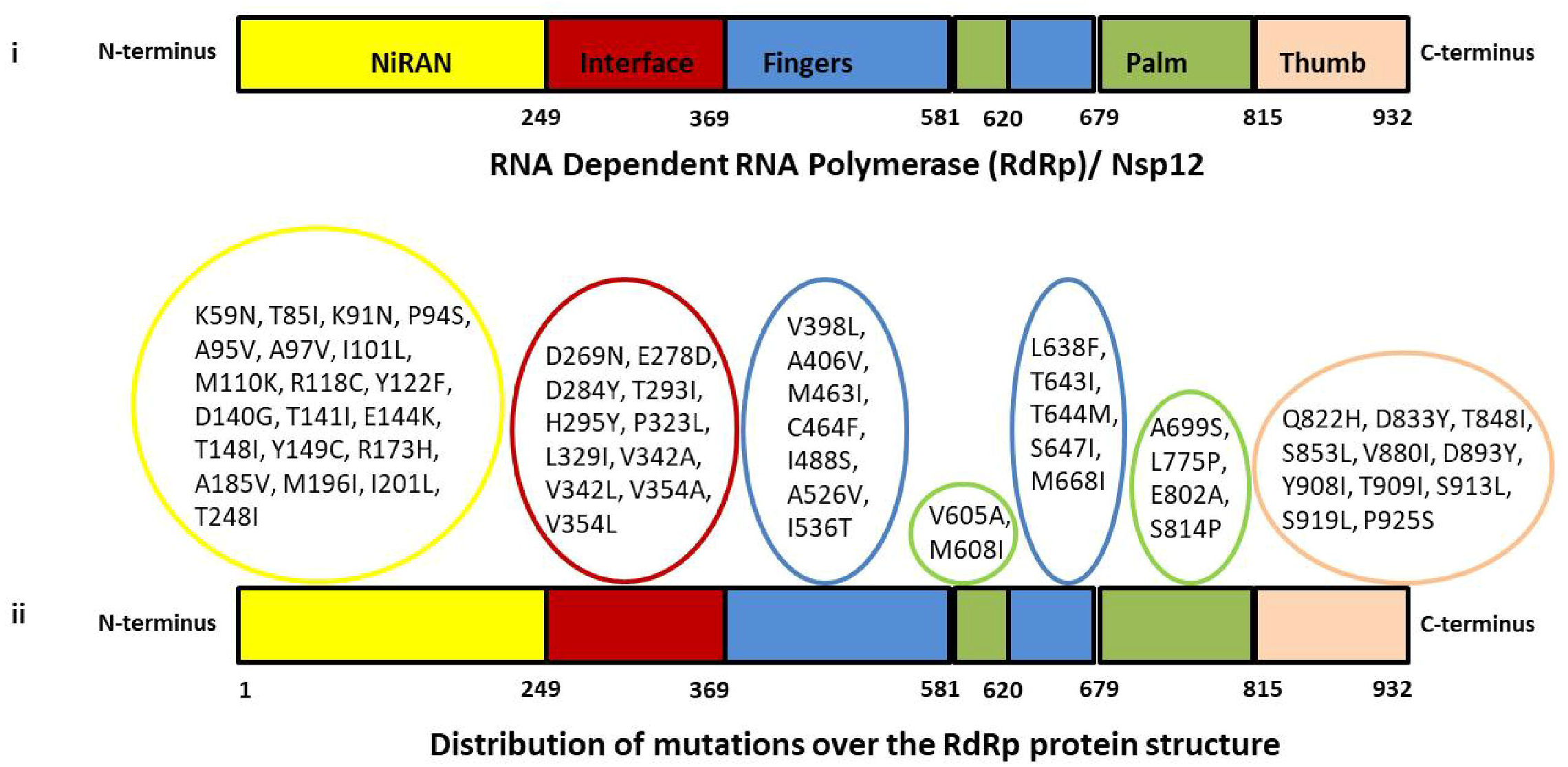
The details of the mutation identified in RdRp among Indian SARS-CoV-2 isolates. A) Schematic diagram of the domain architecture of RdRp. Each domain of RdRp is represented by a unique color. The interdomain borders are labeled with residue numbers. B) The mutations present in RdRp among the Indian isolates of SARS-CoV-2 are demonstrated in the schematics.

### Mutations affect RdRp protein dynamic stability and flexibility

We performed protein modelling studies using DynaMut programe to understand, if the mutation observed in RdRp can alter protein structural integrity. The DynaMut programe does protein modelling for those proteins whose crystal structure has been solved. For this protein modelling study, we used recently reported crystal structure of RdRp (PDB ID: 7BV1)(Yin et al., 2020). First, we calculated the differences in free energy (ΔΔG) between wild-type and mutant. Our data revealed that mutations at twenty-two positions cause stabilisation in protein structure (positive ΔΔG) as shown in table 2, maximum positive ΔΔG was obtained for D833Y (1.372 kcal/mol). Similarly, the mutations at twenty-eight positions cause destabilisation (negative ΔΔG) in protein structure upon mutation (Table 2), maximum negative ΔΔG was obtained for the mutant A699S (−2.233 kcal/mol).

**Table 2.** The ΔΔG and ΔΔSvib ENCOM values obtained by protein modelling using DynaMut programe. The positive and negative ΔΔG represents increase and decrease in protein stability upon mutation. Similarly, the positive and negative ΔΔSvib ENCOM represents the increase in flexibility and rigidity upon mutations.

| S. No. | Mutation | $\Delta\Delta G$<br>(kcal/mol) | $\Delta\Delta S_{vib}$ ENCOM<br>(kcal.mol<sup>-1</sup>.K<sup>-1</sup>) |
| --- | --- | --- | --- |
| 1 | T85I | -0.707 | -0.074 |
| 2 | K91N | -0.311 | 0.295 |
| 3 | P94S | -0.035 | -0.131 |
| 4 | A95V | -0.669 | -0.627 |
| 5 | A97V | 0.469 | -1.020 |
| 6 | R118C | 0.005 | 0.055 |
| 7 | Y122F | 0.225 | 0.273 |
| 8 | D140G | -0.18 | 0.035 |
| 9 | T141I | 1.041 | -0.231 |
| 10 | E144K | -0.236 | 0.353 |
| 11 | T148I | 1.156 | -0.098 |
| 12 | Y149C | 0.237 | 1.065 |
| 13 | R173H | -0.284 | 0.01 |
| 14 | A185V | -0.179 | -0.369 |
| 15 | M196I | -0.554 | 0.094 |
| 16 | I201L | 0.105 | 0.057 |
| 17 | T248I | 0.452 | -0.08 |
| 18 | D269N | -0.362 | 0.08 |
| 19 | E278D | -0.714 | 0.103 |
| 20 | D284Y | 0.346 | -0.459 |
| 21 | T293I | 0.264 | -0.016 |
| 22 | H295Y | 0.304 | 0.141 |
| 23 | P323L | 0.53 | -0.252 |
| 24 | L329I | 0.081 | 0.231 |
| 25 | V342A | -0.454 | 0.402 |
| 26 | V342L | 1.027 | -0.025 |
| 27 | V354A | -1.811 | 0.277 |
| 28 | V354L | -0.162 | -0.387 |
| 29 | V398L | 0.37 | 0.031 |
| 30 | A406V | -0.148 | -0.047 |
| 31 | M463I | 0.483 | 0.25 |
| 32 | C464F | 1.251 | -1.092 |
| 33 | I488S | -1.617 | 0.205 |
| 34 | A526V | 0.305 | -0.109 |
| 35 | I536T | -1.703 | 0.348 |
| 36 | V605A | -1.528 | 0.596 |
| 37 | L638F | -0.167 | -0.297 |
| 38 | T643I | -0.383 | -0.074 |
| 39 | T644M | -0.084 | -0.037 |
| 40 | S647I | -0.272 | 0.166 |
| 41 | M668I | 0.104 | 0.066 |
| 42 | A699S | -2.233 | -0.185 |
| 43 | L775P | -0.6 | 0.754 |
| 44 | E802A | -0.835 | 0.49 |
| 45 | S814P | -0.083 | 0.113 |
| 46 | Q822H | 0.006 | 0.514 |
| 47 | D833Y | 1.372 | -0.397 |
| 48 | V880I | -0.087 | -0.146 |
| 49 | D893Y | 0.088 | -0.168 |
| 50 | S913L | -0.116 | 0.012 |

Next, we measured the changes in vibrational entropy energy (ΔΔSVibENCoM) between the wild type and the mutant. The negative and positive ΔΔSVibENCoM values depict the rigidification and gain in protein flexibility upon mutation, respectively. Our data revealed that mutation at twenty-seven positions causes increase in flexibility of mutant protein (positive ΔΔSVibENCoM). The maximum positive ΔΔSVibENCoM was obtained for Y149C (1.065 kcal.mol-1.K-1) mutant. Similarly, the mutations at rest of the twenty-three positions cause rigidification of protein structure (negative ΔΔSVibENCoM) in protein structure upon mutation (Table 2). The maximum negative ΔΔSVibENCoM was obtained for C464F (−1.092 kcal.mol-1.K-1) mutant. Altogether, our data revealed that the mutation observed in RdRp affects both protein dynamicity and flexibility.

### Identification of B cell epitopes of RdRp

The continuous B-cell epitopes of RdRp were predicted by IEDB webserver tool and the epitopes are shown in figure 2A. The yellow area of the graph corresponds to those regions of the RdRp that can potentially contribute to the B cell epitopes. Our data demonstrated thirty-six epitopes of varying lengths that could potentially act as B cell epitopes (figure 2B). Among those peptides, the ‘peptide 18’ is the largest epitopes, which is forty-four amino acid in length (from RdRp residue 482 to 525). Similarly, peptide 5, 19, 30, 31 and 34 are comprised of single amino acid only (figure 2B).

**Figure 2:**
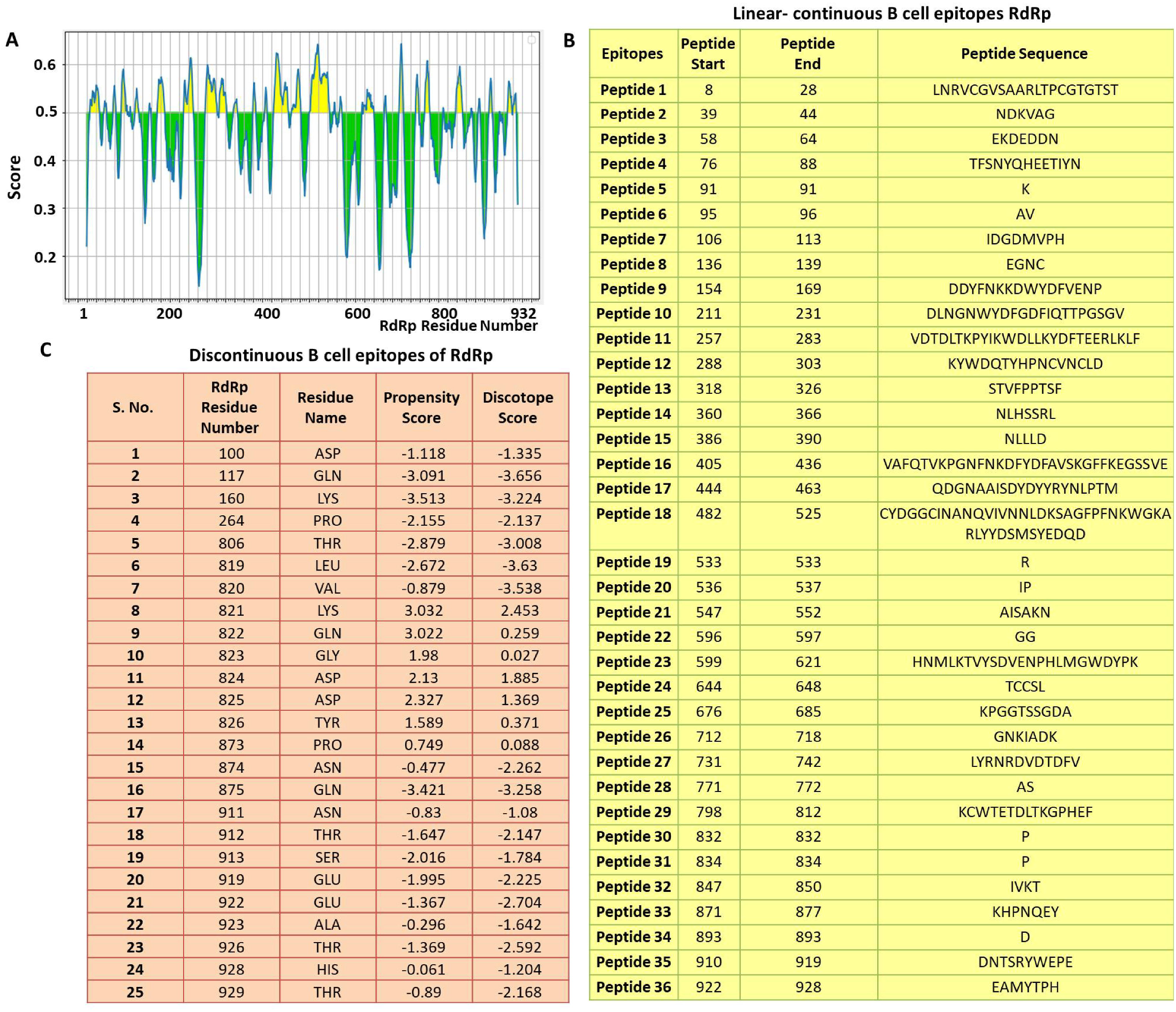
Prediction of B-cell epitopes of RdRp. A) Linear continuous B-cell epitopes contributed by RdRp, the Y-axis of the graph corresponds to BepiPred score, while the X-axis depicts the RdRp residue positions in the sequence. The data was generated by IEDB webserver using ‘Bepipred Linear Epitope Prediction 2.0’ method. The chart is divided into two parts yellow and green. The RdRp residues present in the yellow have higher probability to be part of the linear continuous B cell epitope. B) The details of the linear continuous B cell epitopes are listed. The sequence of each peptide along with its start and end point in the RdRp polypeptide sequence is also mentioned. C) Prediction of discontinuous B-cell epitopes of RdRp by DiscoTope 2.0 web tool. The position of each predicted epitope is mentioned along with its propensity and DiscoTope score.

Subsequently, we predicted the B cell epitopes of RdRp based on its three dimensional structure using DiscoTope 2.0 webserver tool (Kringelum et al., 2012). Our analysis revealed twenty-five discontinuous epitopes of RdRp having high score. The locations of these epitopes are highlighted in figure 2C along with its propensity and DiscoTope score.

Among discontinuous epitopes, approximately 80% of them (20 out of 25) reside towards the C-terminal end of RdRp (from residue 800 to 932) as shown in figure 2C. Altogether, our data revealed B cell epitopes contributed by RdRp.

### RdRp mutants preferentially localises in the B cell epitopes region

Subsequently, we analysed and compared the RdRp mutations that reside in the linear-continuous and discontinuous B cell epitopes. Our data revealed that out of 60 mutants observed in this study, 24 resides in the B cell epitope region of RdRp (figure 3A). These 24 mutants correspond to 40% of the total mutants observed among Indian isolates. The details of all 24 mutants that localises in B cell epitope region are shown in figure 3B. Altogether, our data strongly suggest that RdRp mutants preferentially localises in B cell epitope region.

**Figure 3:**
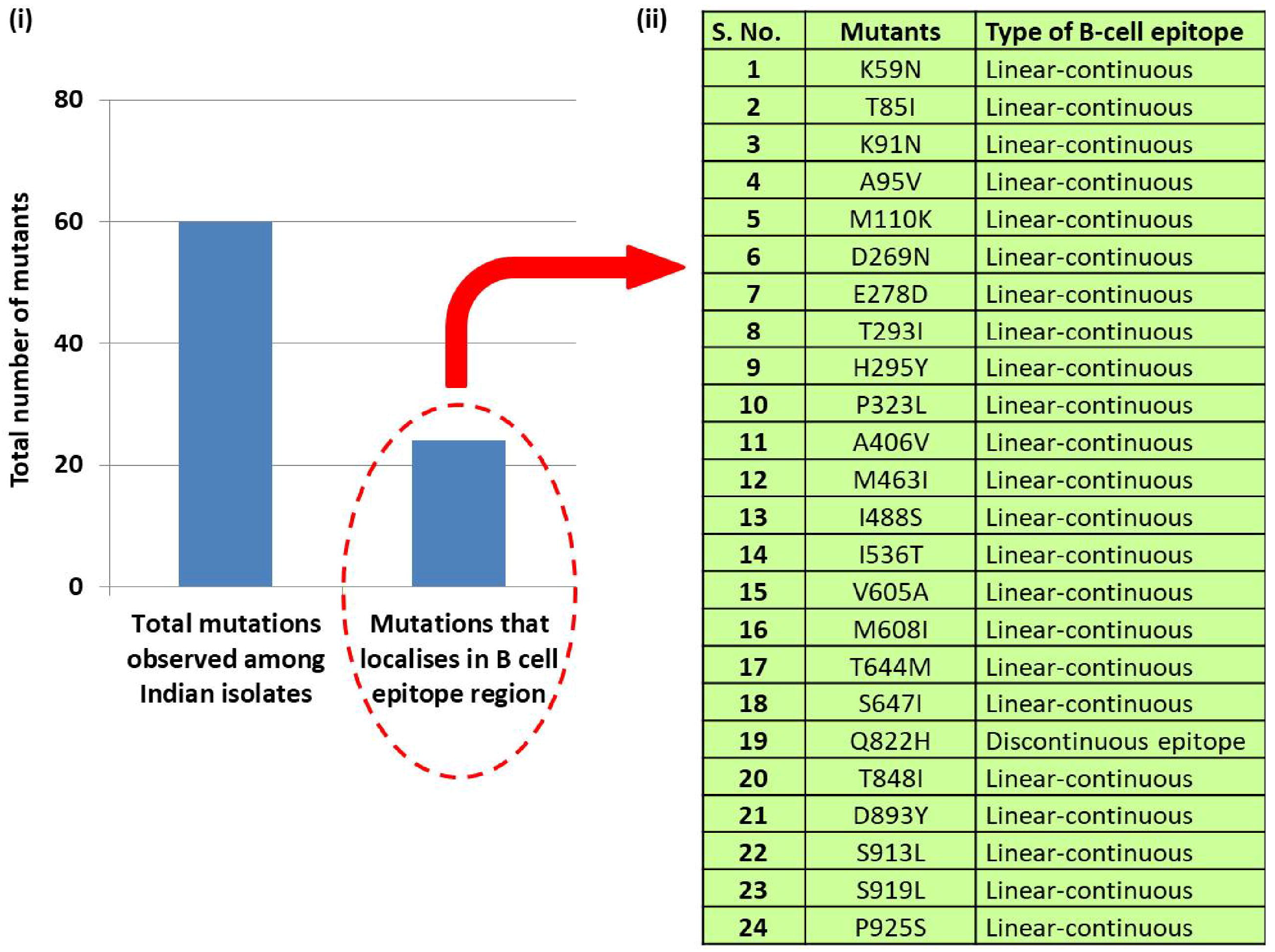
Correlation of RdRp mutants and B cell epitopes. A) the graph shows the distribution of RdRp mutants observed among Indian SARS-CoV-2 isolates. Out of 60, 24 mutants localises in B cell epitope region of RdRp. B) Detail of the RdRp mutants that localises in B cell epitope region.

## DISCUSSION

The coronaviruses belongs to RNA viruses that exhibits high rate of mutations in their genome (Benvenuto et al., 2020). As these viruses spread to new locations they keep on acquiring mutations and few of them are naturally selected because of their beneficial effect on the virus. The SARS-CoV-2 was first reported from Wuhan, china and within a span of few months it spread to almost all countries worldwide (Worldometers, 2020). The researchers around the globe sequenced the SARS-CoV-2 to understand the genomic properties of this virus and soon identify various mutations. The investigation on the genomic variation acquired by SARS-CoV-2 is indispensable for understanding the epidemiology, pathogenesis; devise preventive measures and treatment strategies against COVID-19.

The earlier variation studies on SARS-CoV-2 revealed that RdRp is among the mutational hotspot protein (Pachetti et al., 2020). In the similar directions, this study was conducted with an aim to identify mutations in RdRp from Indian isolates. Our earlier study revealed seven crucial mutations in RdRp of SARS-CoV-2 (Chand et al., 2020) that can have potential impact on this protein function. The present study identifies and characterises B cell epitope contributed by RdRp and correlate them with the observed mutants. In this study, we analysed 668 RdRp sequences reported from India till April 2021 date and identified sixty mutations in RdRp. Our protein modelling studies revealed various interesting mutations including D833Y, A699S, Y149C and C464F (table 2) that can potentially affect stability and flexibility of RdRp. So far, we analysed 668 sequences from India and identified sixty mutations which indicates that RdRp is one of the mutational hotspot protein of SARS-CoV-2 in India. Furthermore, we did prediction study to identify B cell epitopes contributed by RdRp. Our data revealed that there are thirty-six high rank linear-continuous B cell epitope as well as twenty-five discontinuous B cell epitopes. Moreover, we also identified that out of sixty mutants identified among Indian isolates, twenty-four resides (40%) in these B cell epitope region.

The variations in RdRp or any other protein of SARS-CoV-2 will possibly tell us how the virus is evolving. Earlier studies with RNA viruses have also shown that these viruses keep on mutating to better adapt and survive in the host (Sanjuán and Domingo-Calap, 2016). Here, in this study, we have reported RdRp mutations, its correlation with B cell epitopes. However, it warrants future studies to understand the possible effect of these mutations on virus infectivity and life cycle.

## CONCLUSIONS

The pandemic caused by SARS-CoV-2 have adverse impact on health services worldwide with economic fallout of most of the countries. Understanding the genetic variations in SARS-CoV-2 will help to better devise the preventive and treatment strategies against this virus. Here, our data show various mutations in RdRp from India and their impact on B cell epitopes by *in silico* studies. Altogether, the data presented here could help scientific community to better understand the immunological aspect of SARS-CoV-2.

## ACKNOWLEDGEMENTS

We would like to acknowledge Patna University, Patna, Bihar (India) for providing infrastructural support for this study.

---

**Supplementary Table 1:** List of protein identifier accession number used in this analysis. The sequences of each protein can be downloaded from NCBI-Virus-Database using the accession number provided in the list.

|  |  |  |  |  |  |  |
| --- | --- | --- | --- | --- | --- | --- |
| YP_009724389 | QOS50986.1 | QNL90834.1 | QLI49743.1 | QKY60095.1 | QKJ84977.1 | QJX44644.1 |
| QTV76173.1 | QOS51010.1 | QNL90846.1 | QLI49755.1 | QKY60107.1 | QKG91164.1 | QJX44656.1 |
| QTP17383.1 | QOS51022.1 | QNL90858.1 | QLI49767.1 | QKY60119.1 | QKG91176.1 | QJX44668.1 |
| QTH36343.1 | QOS51058.1 | QNL90870.1 | QLI49779.1 | QKY60131.1 | QKG91188.1 | QJX44680.1 |
| QTH36355.1 | QOS51070.1 | QNL90882.1 | QLI49791.1 | QKY60163.1 | QKG91200.1 | QJW39842.1 |
| QTH36367.1 | QOS51082.1 | QNL90894.1 | QLI49803.1 | QKY60175.1 | QKG91212.1 | QJW39854.1 |
| QTH36379.1 | QOS51094.1 | QNL90906.1 | QLI49815.1 | QKY60187.1 | QKG91224.1 | QJW39866.1 |
| QTH36391.1 | QOS51106.1 | QNL90918.1 | QLI52055.1 | QKY60199.1 | QKG91236.1 | QJW39878.1 |
| QTH36403.1 | QOS51118.1 | QNL98443.1 | QLI52067.1 | QKY60211.1 | QKG91248.1 | QJW39890.1 |
| QTH36415.1 | QOR63432.1 | QNL98455.1 | QLH64777.1 | QKY60223.1 | QKG91260.1 | QJW39902.1 |
| QTH36427.1 | QOR63444.1 | QNL98467.1 | QLH64789.1 | QKY60235.1 | QKG91272.1 | QJW39914.1 |
| QTH36439.1 | QOR63456.1 | QNL98479.1 | QLH64801.1 | QKY60262.1 | QKG91284.1 | QJW39926.1 |
| QTH36451.1 | QOR63468.1 | QNL35816.1 | QLH64813.1 | QKY64328.1 | QKH78808.1 | QJW69137.1 |
| QTH36723.1 | QOR63480.1 | QNL35828.1 | QLH64825.1 | QKY64344.1 | QKH78820.1 | QJU70555.1 |
| QTG68689.1 | QOR63504.1 | QNL35840.1 | QLH64837.1 | QKY64358.1 | QKI10470.1 | QJT43414.1 |
| QTG68602.1 | QOR64231.1 | QNL35852.1 | QLH64849.1 | QKY64612.1 | QKI28575.1 | QJT43426.1 |
| QTG68615.1 | QOR64243.1 | QNL35864.1 | QLH64861.1 | QKY64638.1 | QKI28587.1 | QJT43438.1 |
| QTG40603.1 | QOQ57010.1 | QNL35876.1 | QLH64875.1 | QKY64790.1 | QKI28599.1 | QJT43450.1 |
| QRY06669.1 | QOQ57022.1 | QNL35888.1 | QLH64887.1 | QKY65275.1 | QKI28611.1 | QJT43462.1 |
| QRQ47003.1 | QOQ57034.1 | QNL35900.1 | QLH64899.1 | QKV25883.1 | QKI28623.1 | QJT43474.1 |
| QRQ47018.1 | QOQ57046.1 | QNL35912.1 | QLH64915.1 | QKV25895.1 | QKI28635.1 | QJT43486.1 |
| QRQ47032.1 | QOQ57058.1 | QNL35924.1 | QLH64927.1 | QKV25907.1 | QKI28647.1 | QJT43498.1 |
| QRQ47071.1 | QOQ57070.1 | QNL35936.1 | QLH64939.1 | QKV25919.1 | QKI28659.1 | QJT43510.1 |
| QRQ47083.1 | QOQ57082.1 | QNL35948.1 | QLH90040.1 | QKV25931.1 | QKI28671.1 | QJT43522.1 |
| QRQ69195.1 | QOQ57094.1 | QNL35960.1 | QLH90076.1 | QKV25943.1 | QKI28683.1 | QJT43534.1 |
| QRQ69279.1 | QOQ57106.1 | QNL35972.1 | QLH90088.1 | QKV25955.1 | QKE61658.1 | QJT43546.1 |
| QRN68229.1 | QOQ57119.1 | QNL35984.1 | QLH90101.1 | QKV25967.1 | QKE61670.1 | QJT43558.1 |
| QRN68241.1 | QOQ72542.1 | QNL35996.1 | QLH93117.1 | QKV25979.1 | QKE61682.1 | QJT43570.1 |
| QRN68253.1 | QOQ72554.1 | QNL36008.1 | QLH93129.1 | QKV25991.1 | QKE61694.1 | QJT43582.1 |
| QRN68266.1 | QOQ72566.1 | QNL36560.1 | QLH93185.1 | QKV26003.1 | QKE61706.1 | QJT43594.1 |
| QRN68278.1 | QOQ72578.1 | QLY82444.1 | QLH93199.1 | QKV26015.1 | QKE61718.1 | QJT43606.1 |
| QRN68302.1 | QOQ84793.1 | QLY82482.1 | QLH93212.1 | QKV26027.1 | QKE61730.1 | QJT43618.1 |
| QRN68400.1 | QOQ84824.1 | QLY82500.1 | QLH93283.1 | QKV26039.1 | QKE61742.1 | QJT43630.1 |
| QRN68424.1 | QKM77202.1 | QLY82512.1 | QLF97937.1 | QKV26051.1 | QKE61754.1 | QJT43642.1 |
| QRN68442.1 | QKM77214.1 | QMC85346.1 | QLF97949.1 | QKV26063.1 | QKE61766.1 | QJT43654.1 |
| QRM36235.1 | QKM77226.1 | QLQ87394.1 | QLF97985.1 | QKV26075.1 | QKE61778.1 | QJT43666.1 |
| QRM91443.1 | QKM77238.1 | QLQ87406.1 | QLF97997.1 | QKV26087.1 | QKE61790.1 | QJT43678.1 |
| QRE02629.2 | QKM77250.1 | QLQ87418.1 | QLF98069.1 | QKV27549.1 | QKE61802.1 | QJT43690.1 |
| QRC50281.1 | QKM77262.1 | QLQ87430.1 | QLF98081.1 | QKV27561.1 | QJY77053.1 | QJT43702.1 |
| QRC50293.1 | QKM77274.1 | QLQ87442.1 | QLF98093.1 | QKV27573.1 | QJY51262.1 | QJT43714.1 |
| QRC50305.1 | QNO30921.1 | QLQ87454.1 | QLF98105.1 | QKV27585.1 | QJY51274.1 | QJT43726.1 |
| QRC50340.1 | QNO30933.1 | QLQ87466.1 | QLF98117.1 | QKQ29884.1 | QJY51286.1 | QJS39637.1 |
| QRC50364.1 | QNN87972.1 | QLQ87478.1 | QLF98129.1 | QKQ29944.1 | QJY51346.1 | QJS39649.1 |
| QRC50492.1 | QNN87984.1 | QLQ87490.1 | QLF98141.1 | QKQ29956.1 | QJY51358.1 | QJR84343.1 |
| QQJ94103.1 | QNN87996.1 | QLQ87502.1 | QLF98153.1 | QKQ29968.1 | QJY51370.1 | QJR84367.1 |
| QQJ95306.1 | QNN88008.1 | QLQ87514.1 | QLF98165.1 | QKQ30028.1 | QJY51382.1 | QJR84379.1 |
| QQH18637.1 | QNN88020.1 | QLQ87526.1 | QLF98178.1 | QKQ30040.1 | QJY40383.1 | QJR84391.1 |
| QPZ33508.1 | QNN88068.1 | QLQ87550.1 | QLF98198.1 | QKQ30052.1 | QJY40395.1 | QJR84415.1 |
| QPB40017.1 | QNN88080.1 | QLQ87574.1 | QLF98210.1 | QKQ30064.1 | QJY40407.1 | QJR84427.1 |
| QPB40029.1 | QNN88092.1 | QLQ87598.1 | QLF98222.1 | QKQ30076.1 | QJY40419.1 | QJR84439.1 |
| QPB40041.1 | QNN88104.1 | QLQ87610.1 | QLF98234.1 | QKQ30088.1 | QJY40431.1 | QJR84451.1 |
| QPB40053.1 | QNN88116.1 | QLQ87622.1 | QLF98246.1 | QKQ30100.1 | QJY40443.1 | QJR84463.1 |
| QPB40065.1 | QNN88128.1 | QLQ87634.1 | QLF98258.1 | QKQ30112.1 | QJY40455.1 | QJR84475.1 |
| QPB40077.1 | QNN88140.1 | QLQ87658.1 | QLF98275.1 | QKQ30124.1 | QJY40467.1 | QJR84487.1 |
| QPB40089.1 | QNN88152.1 | QLQ87682.1 | QLF98287.1 | QKQ30136.1 | QJY40479.1 | QJR84499.1 |
| QPB40101.1 | QNN88164.1 | QLQ87694.1 | QLA09686.1 | QKQ30148.1 | QJY40491.1 | QJR84511.1 |
| QPB40113.1 | QNN88176.1 | QLQ87706.1 | QLA09698.1 | QKQ30160.1 | QJY40503.1 | QJR84523.1 |
| QPB40563.1 | QNN88188.1 | QLQ87718.1 | QLA09710.1 | QKQ30172.1 | QJY40515.1 | QJR84535.1 |
| QPB40575.1 | QNN88200.1 | QLQ87730.1 | QLA09722.1 | QKQ30184.1 | QJY40527.1 | QJQ28343.1 |
| QPB17918.1 | QNN88212.1 | QLQ87795.1 | QLA09734.1 | QKQ30196.1 | QJY40539.1 | QJQ28355.1 |
| QPB17990.1 | QNN88224.1 | QLQ87983.1 | QLA09746.1 | QKQ30208.1 | QJY40551.1 | QJQ28367.1 |
| QPB18014.1 | QNN88236.1 | QLQ87995.1 | QLA09758.1 | QKQ30220.1 | QJY40563.1 | QJQ28379.1 |
| QPB18026.1 | QNN88248.1 | QLQ88046.1 | QLA09770.1 | QKQ30232.1 | QJY40575.1 | QJQ28391.1 |
| QPB18050.1 | QNN88260.1 | QLQ88063.1 | QLA09782.1 | QKQ30244.1 | QJY40587.1 | QJQ28403.1 |
| QPB18062.1 | QNN88272.1 | QLQ88075.1 | QLA09794.1 | QKQ30256.1 | QJY40599.1 | QJQ28415.1 |
| QPB18074.1 | QNN88284.1 | QLQ88087.1 | QLA09808.1 | QKQ30268.1 | QJW00289.1 | QJQ28427.1 |
| QPB18086.1 | QNN88296.1 | QLQ88102.1 | QLA09820.1 | QKQ63386.1 | QJW00301.1 | QJH92165.1 |
| QPB18098.1 | QNN88551.1 | QLR06736.1 | QLA09832.1 | QKO00484.1 | QJW00313.1 | QJH92177.1 |
| QPB18110.1 | QNN88563.1 | QLR06757.1 | QLA09844.1 | QKJ68410.1 | QJW00325.1 | QJF77844.1 |
| QPB18122.1 | QNN88575.1 | QLR06769.1 | QLA09856.1 | QKJ68422.1 | QJW00337.1 | QJF77856.1 |
| QPB18134.1 | QNN88630.1 | QLR06870.1 | QLA09880.1 | QKJ68434.1 | QJW00349.1 | QJF77868.1 |
| QPB18158.1 | QNN88642.1 | QLR06882.1 | QLA09904.1 | QKJ68446.1 | QJW00361.1 | QJF77880.1 |
| QPB18194.1 | QNN88654.1 | QLR06894.1 | QLA10066.1 | QKJ68458.1 | QJW00373.1 | QJC19489.1 |
| QPB18206.1 | QNN90122.1 | QLR07146.1 | QLA10078.1 | QKJ68471.1 | QJW00385.1 | QHS34545.1 |
| QPB39980.1 | QNN90135.1 | QLR07158.1 | QLA10090.1 | QKJ68483.1 | QJW00397.1 | QIA98582.1 |
| QPB39992.1 | QNN93013.1 | QLR07170.1 | QLA10102.1 | QKJ68495.1 | QJW00409.1 |  |
| QPB40004.1 | QNN26334.1 | QLR07182.1 | QLA10114.1 | QKJ68507.1 | QJW00421.1 |  |
| QPA18453.1 | QNN26346.1 | QLR07194.1 | QLA10126.1 | QKJ68519.1 | QJW00433.1 |  |
| QPA18457.1 | QNN26358.1 | QLR07206.1 | QLA10138.1 | QKJ68531.1 | QJW00445.1 |  |
| QOW08181.2 | QNN26370.1 | QLR12235.1 | QLA10150.1 | QKJ68543.1 | QJW00457.1 |  |
| QOU99168.1 | QNN26382.1 | QLR12247.1 | QLA10162.1 | QKJ68555.1 | QJX44380.1 |  |
| QOU99261.1 | QNN26394.1 | QLR12259.1 | QLA10174.1 | QKJ68567.1 | QJX44392.1 |  |
| QOU99272.1 | QNN26406.1 | QLR12271.1 | QLA10186.1 | QKJ68579.1 | QJX44404.1 |  |
| QOU99283.1 | QNN26418.1 | QLR12283.1 | QLA10198.1 | QKJ68591.1 | QJX44416.1 |  |
| QOU99294.1 | QNN26430.1 | QLR12295.1 | QLA10210.1 | QKJ68603.1 | QJX44428.1 |  |
| QOS50509.1 | QNN26442.1 | QLR12307.1 | QLA10222.1 | QKJ68615.1 | QJX44440.1 |  |
| QOS50580.1 | QNN26454.1 | QLR12319.1 | QKY74626.1 | QKJ68627.1 | QJX44464.1 |  |
| QOS50592.1 | QNN26466.1 | QLR12331.1 | QKY74638.1 | QKJ68639.1 | QJX44476.1 |  |
| QOS50604.1 | QNN26478.1 | QLR12343.1 | QKY59939.1 | QKJ68651.1 | QJX44500.1 |  |
| QOS50616.1 | QNN30834.1 | QLR12355.1 | QKY59951.1 | QKJ68663.1 | QJX44512.1 |  |
| QOS50628.1 | QNN30858.1 | QLR12415.1 | QKY59963.1 | QKJ68675.1 | QJX44536.1 |  |
| QOS50676.1 | QNN30870.1 | QLR12438.1 | QKY59975.1 | QKJ68687.1 | QJX44548.1 |  |
| QOS50712.1 | QNN31194.1 | QLR60574.1 | QKY59987.1 | QKJ68699.1 | QJX44560.1 |  |
| QOS50879.1 | QNN31206.1 | QLR80398.1 | QKY59999.1 | QKJ68711.1 | QJX44572.1 |  |
| QOS50891.1 | QNN81330.1 | QLS04049.1 | QKY60023.1 | QKJ68723.1 | QJX44584.1 |  |
| QOS50926.1 | QNN81490.1 | QLI49695.1 | QKY60047.1 | QKJ68735.1 | QJX44596.1 |  |
| QOS50938.1 | QNN81502.1 | QLI49707.1 | QKY60059.1 | QKJ84941.1 | QJX44608.1 |  |
| QOS50950.1 | QNN83660.1 | QLI49719.1 | QKY60071.1 | QKJ84953.1 | QJX44620.1 |  |
| QOS50962.1 | QNN83672.1 | QLI49731.1 | QKY60083.1 | QKJ84965.1 | QJX44632.1 |  |

